# TatA and TatB generate a hydrophobic mismatch that is important for function and assembly of the Tat translocon in *Escherichia coli*

**DOI:** 10.1101/2021.05.26.445790

**Authors:** Denise Mehner-Breitfeld, Michael T. Ringel, Daniel Alexander Tichy, Laura J. Endter, Kai Steffen Stroh, Heinrich Lünsdorf, Herre Jelger Risselada, Thomas Brüser

## Abstract

The Tat system has the unique purpose to translocate folded proteins across energy-transducing membranes. It occurs in bacteria and archaea, as well as in eukaryotic organelles of bacterial origin. In the bacterial model system *Escherichia coli*, the three components TatA, TatB, and TatC assemble to functional translocons. TatA and TatB both possess an N-terminal transmembrane helix (TMH) that is followed by an amphipathic helix (APH). The TMHs of TatA and TatB generate a hydrophobic mismatch with only 12 consecutive hydrophobic residues that span the membrane. We shortened or extended this stretch of hydrophobic residues in either TatA, TatB, or both, and analyzed effects on transport functionality and translocon assembly. The wild type length functioned best but was not an absolute requirement, as some variation was clearly tolerated. Defects of shortenings or extensions were enhanced by simultaneous mutations in TatA and TatB, indicating partial compensations of mutations in TatA by wild type TatB or vice versa. Length variation in TatB destabilized TatBC-containing complexes, revealing that the 12-residues-length is important for Tat component interactions and translocon assembly. To also address potential effects on TatA associations, we characterized these by metal tagging transmission electron microscopy and carried out molecular dynamics simulations. In these simulations, interacting short TMHs of larger TatA assemblies were thinning the membrane together with laterally aligned tilted APHs that generated a deep V-shaped groove. The conserved length of 12 hydrophobic residues may thus not only be important for translocon interactions, but also for a membrane destabilization during Tat transport. If this is the case, the specific short length could be a compromise between functionality and proton leakage minimization.

## Introduction

Tat systems serve the purpose to transport folded proteins across energy-transducing membranes in bacteria, archaea, plastids, and some mitochondria [1–3]. They minimally consist of two components, TatA and TatC [4], but three-component TatABC systems are very common and found in the model Tat systems of *Escherichia coli* and plant plastids [5]. TatA and TatB are structurally similar and evolutionary related [6]. Both are membrane-anchored by a 13-15 residues long N-terminal transmembrane helix (TMH) that contains only 12 consecutive hydrophobic residues. This helix is connected to an amphipathic helix (APH) via a short hinge at the cytoplasmic surface of the membrane [7–10] (Fig 1A). The APH is followed by a negatively charged patch of residues in TatA [11], and more C-terminal regions are neither conserved nor functionally essential [12]. TatC has six transmembrane domains with both termini on the cytoplasmic side [13,14]. TatB interacts tightly with TatC, whereas TatA gradually dissociates during purifications [15–17]. From studies employing cross-linking and fluorescent protein tagging, it is known that the TatA interaction with TatBC changes during the translocation cycle, and larger TatA assemblies at TatABC complexes are believed to enable transport [18–23]. A subpopulation of TatA protomers interacts with TatBC in a way that is not affected by mutations in TatC that abolish substrate-binding [15], and resting-state-contacts of TatA to TatC have been identified [24]. TatA assemblies undergo substrate-induced conformational changes that relate to Tat transport [25,26], and the TatABC components rearrange during the translocation cycle [27,28].

**Fig 1:**
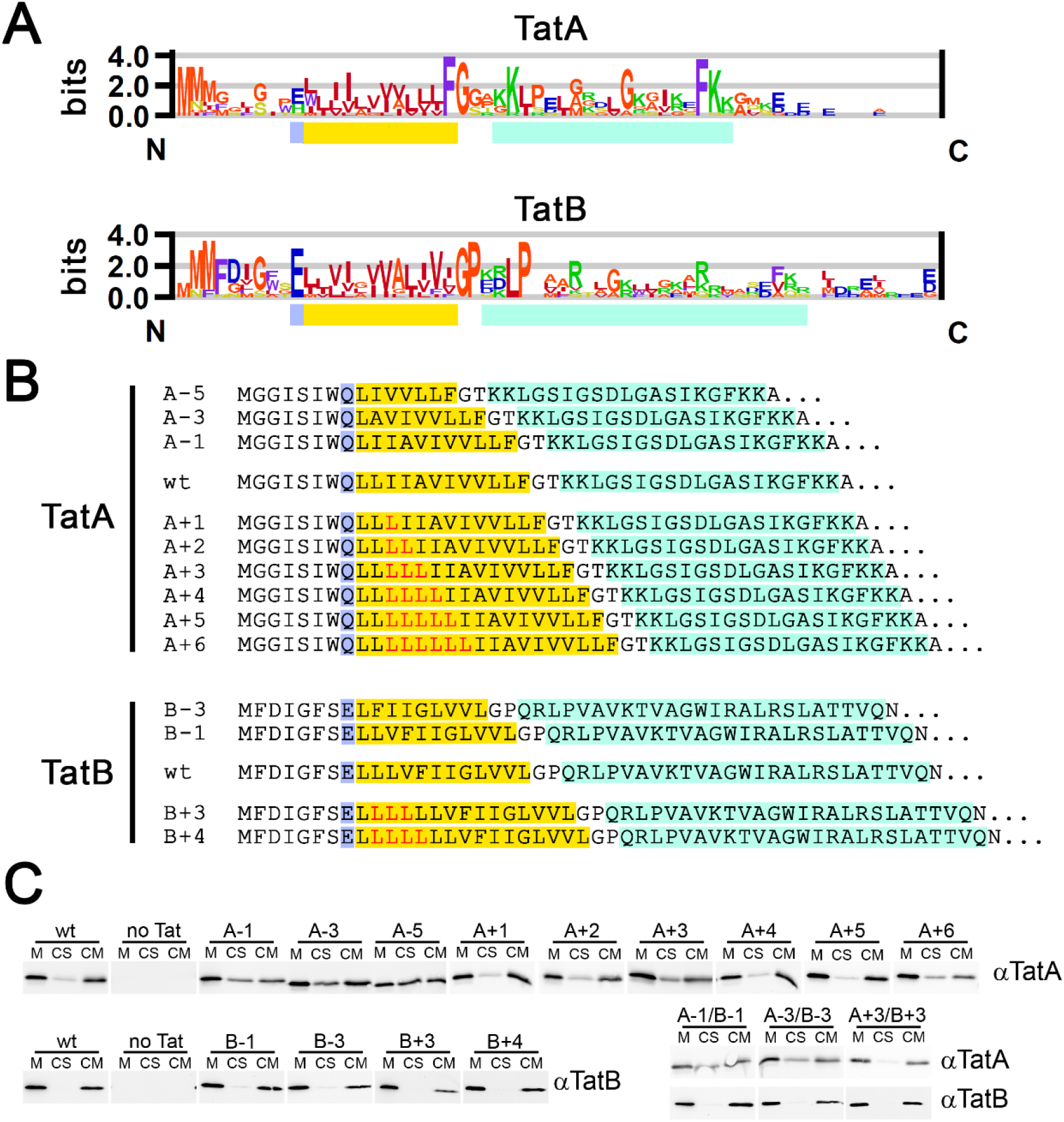
Constructs for the analysis of the 12-residues hydrophobic stretch in TatA and TatB. (A) Sequence logos of the functionally important region of TatA and TatB. Highlighted are the 12-residues hydrophobic stretch in the transmembrane helix of TatA and TatB (yellow), the preceding polar position (bluish purple), and the amphipathic helix (turquoise). The sequence logos have been obtained using enoLOGOS [29] based on sequence alignments by Clustal Omega [30] of sequences from a wide range of organisms, including α-, β-, γ-, *δ*-, and ε-proteobacteria, firmicutes, actinobacteria, cyanobacteria, chlorobi, chloroflexaceae, thermodesulfobacteria, aquificae, and archaea. (B) TatA and TatB variants used in this study. The color code is the same as in (A). Inserted Leu residues are in red. (C) Carbonate wash analysis for the assessment of membrane integration of the TatA and TatB variants used in this study. TatA or TatB variants were detected by SDS-PAGE/Western blotting in the untreated membrane fraction (M), supernatant after carbonate wash (CS), and membranes after carbonate wash (CM). Note that all analyzed TatA and TatB variants are membrane inserted, but TatB variants are more stably membrane inserted than TatA variants, most likely due to their tighter interaction with TatC.

Tat-dependently transported proteins possess an N-terminal signal peptide that contains a highly conserved eponymous twin-arginine motif [31,1]. This motif is recognized by TatBC complexes that catalyze the membrane insertion of the signal peptide [32], whereas TatA is primarily required for the translocation step [18]. TatA shows a high tendency to self-associate [33–36], but high-resolution structures are only available for detergent-solubilized TatA [7,8,10], and the mode of TatA self-interactions in membranes is still unclear. Spin-labeling studies indicated that the transmembrane helices of TatA laterally interact [37], but only circular arrangements of aligned TatA have been suggested [8], and the orientation of the APH has not been included in the latest models [38]. Interestingly, a dimeric structure of detergent-solubilized TatA indicated that also APHs of neighboring TatA protomers laterally interact [10].

TMHs need to span the ca. 3 nm thick hydrocarbon core of membranes, and about 20 hydrophobic residues could in principle do this, but TMHs are in average 24.0 +/- 5.6 residues long [39], which is generally due to flexing or tilting of helices in membranes [40]. It is therefore very unusual that TatA and TatB have membrane anchors with TMHs of 13-15 residues that contain only 12 consecutive hydrophobic residues, and that the length of this 12-residues stretch is strictly conserved from archaea to bacteria (Fig 1A). As this short length does not suffice to span a membrane lipid bilayer of normal thickness, the TMHs of TatA and TatB generate a hydrophobic mismatch. It has been found that the TMH of TatA per se can destabilize membranes [25], but potential effects of helix shortenings or extensions have never been systematically analyzed. We therefore changed the hydrophobic mismatch of TatA and TatB by generating helix shortenings and extensions, and investigated effects on Tat functionality and translocon assembly. Results indicate that both components tolerate smaller shortenings or extensions, but the natural 12 consecutive hydrophobic residues appear to function best. We demonstrate that, in case of TatB, this hydrophobic mismatch is important for the stable interaction of TatB with TatC, and therefore the short helices most likely trigger also the association of TatA with TatBC. To also analyze potential effects on the so far uncharacterized larger TatA assemblies that are present at Tat translocons during the translocation cycle, we visualized such assemblies in whole cells by metal-tagging transmission electron microscopy (METTEM), and carried out MD simulations. As a result of these simulations, we found that the short length of the TMH, together with the APH, has the potential to locally thin and destabilize the membrane. As such an effect could be the basis for membrane permeabilization by Tat systems, which would require a delicate compromise between membrane stability and permeabilization, this might explain the strict conservation of the short length of the TMHs in TatA, TatB, and the TatAB-interacting helices of TatC.

## Results

### Tat systems tolerate some shortening and extension of the stretch of 12 consecutive hydrophobic residues in the TMHs of TatA and TatB

To analyze the potential role of the short stretch of 12 consecutive hydrophobic residues in the TMHs of TatA and TatB, we shortened the hydrophobic helix by removing 1, 3, or 5 residues in TatA, and 1 or 3 residues in TatB. We also extended the helix by 1, 2, 3, 4, 5, or 6 leucine residues in TatA, and by 3 or 4 leucine residues in TatB (Fig 1B). In the following, the nomenclature of the constructs is facilitated for the reader, with “A+1” standing for the construct with one Leu inserted into the helix of TatA, and “B-3” standing for the construct with three hydrophobic residues deleted in the helix of TatB, as examples. To ensure the correct ratio of the Tat proteins, all genetic constructs were generated with the *tatABC* operon under control of its natural promoter in the Tat complementation vector pABS-*tatABC* [41]. We used the *tat*-deletion strain DADE [42] in combination with these vectors, which permitted functional analyses as well as BN-PAGE detections of the Tat complexes. All TatA and TatB variants were stably produced, and carbonate washes revealed that all of them were membrane-integrated (Fig 1C). A significant portion of all TatA variants was released from the membranes, which is already known for TatA and may be due to the presence of these proteins in destabilized membrane regions [25]. The amount of TatA that was released by carbonate washes was higher in case of shortened TMHs (A-1, A-3, and A-5, Fig 1C). As TatA can destabilize membranes, these results may indicate that shorter membrane anchors enhance the membrane-destabilizing effect of TatA. TatB and its variants were more stably membrane-integrated, which is expected as TatB tightly associates with TatC. We then analyzed Tat functionality with the described Tat systems. As Tat-deficient strains become SDS-sensitive, which relates to cell wall defects due to Tat-requirement for transport of the cell wall amidases AmiA and AmiC [43], we assessed Tat functionality by monitoring SDS resistance of the respective strains (Fig 2A).

**Fig 2:**
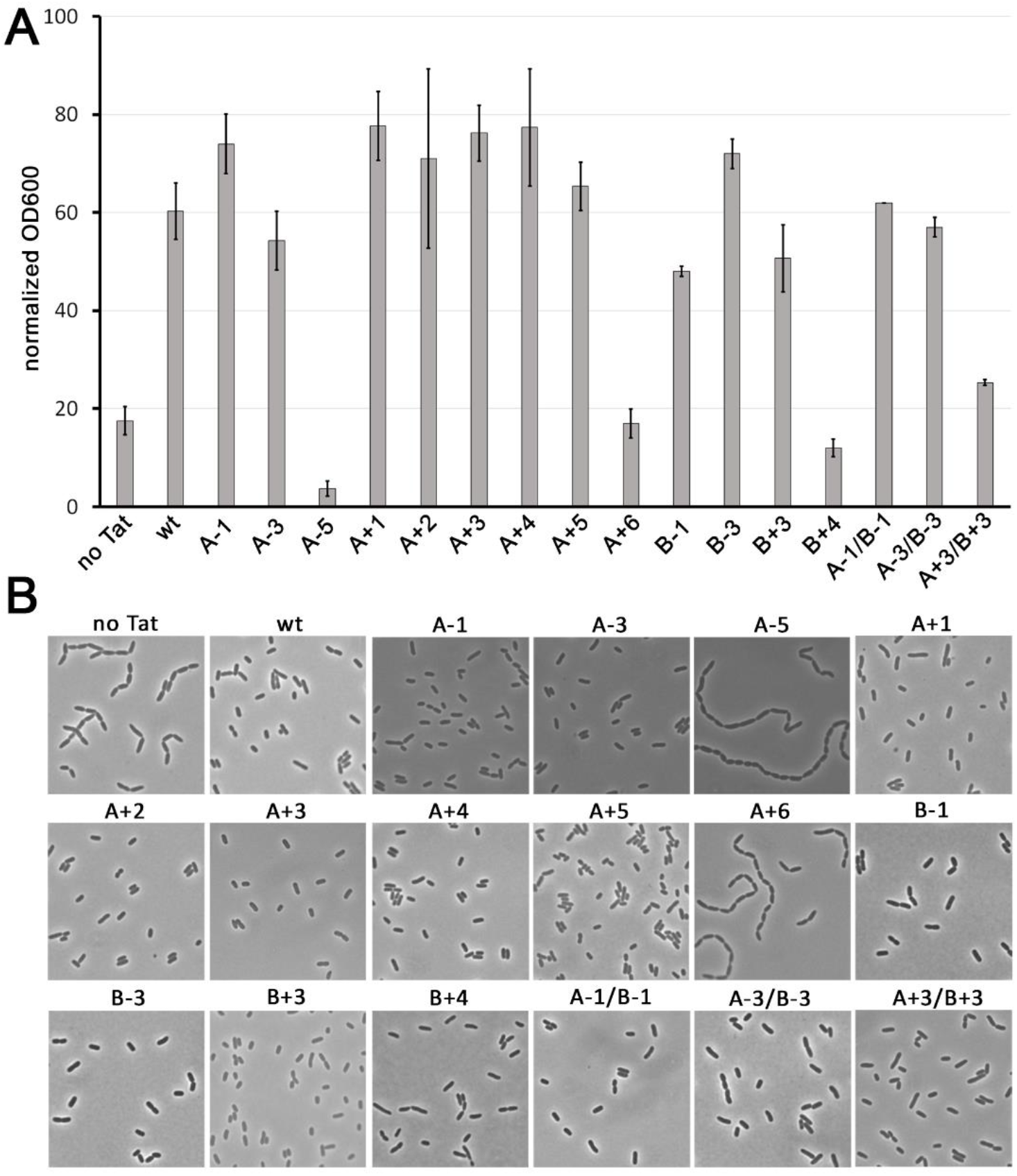
Physiological Tat functionality assays with mutated Tat systems. Complementation of SDS sensitivity (A) and chain formation (B) phenotypes of the Tat deficient strain DADE by Tat systems with indicated single or combined TatA and TatB variants. Controls: ‘no Tat’, empty vector control; ‘wt’, wild type Tat system.

Only the A-5 construct was inactive among the TatA variants with shortenings, and only the A+6 construct was inactive among the extension variants. The A-5 deletion variant showed higher sensitivity towards SDS then the non-complemented *tat* deletion strain, indicating that the A-5 variant was not only non-functional but also had an additional negative physiological effect.

In case of TatB, the B-1 and B-3 deletion-variants were both active in this assay, indicating that shorter hydrophobic helices in principle suffice to establish functional Tat translocons. This was unexpected as TatB tightly interacts with TatC for function, and we therefore expected less tolerance. The Tat system also tolerated the 3-residues extension in TatB, but not anymore the 4-residues extension. As TatA and TatB possibly could have overlapping functions and therefore could complement defects of each other, we also tested combinations of single and triple deletions in TatA and TatB, and triple extensions in TatA and TatB. Notably, the combined deletions were functional, indicating that indeed the deletions did not inactivate the proteins. The combined extension (A+3/B+3) showed a markedly reduced SDS resistance, indicating that some functional overlap had masked the effect of single +3 extensions. Nevertheless, the partial SDS resistance proofed residual activity of A+3 and B+3 variants.

A second way to monitor functionality is the chain formation phenotype, which is similarly based on the absence of AmiA and AmiC in the periplasm of Tat-deficient strains. If these amidases are not transported into the periplasm, the murein is not efficiently hydrolyzed between separating cells, resulting in chains of cells. In full agreement with the SDS-sensitivity measurements, the chain formation phenotype was only observed in those strains that were also SDS-sensitive (Fig 2B). The strain with the A+3/B+3 combination, which had reduced but not abolished SDS resistance, showed no chain formation phenotype, indicating that sufficient amidases were transported for cell separation and confirming the residual activity of the Tat system with the +3 extensions in TatA and TatB. In line with that, the SDS-sensitive B+4 strain showed no obvious chain formation, indicating that transported amidases sufficed for cell separation but not for SDS resistance.

To recognize weaker effects on protein transport, we then carried out biochemical analyses of the transport of the Tat model substrate HiPIP, which is an iron-sulfur cluster containing protein that strictly requires the Tat system for transport [15]. Subcellular fractions were prepared from exponentially growing cells and analyzed by SDS-PAGE/Western blotting using antibodies specifically recognizing HiPIP (Fig 3). We used a vector for low-level constitutive HiPIP production, which results in complete translocation of HiPIP into the periplasm in case of fully functional Tat systems [44]. In this assay, the mature periplasmic HiPIP band indicates transport, and any precursor in the cytoplasm indicates reduced translocation relative to the fully functional Tat system. The wild type Tat system, which served as positive control, showed a strong band of transported mature HiPIP in the periplasm and no accumulation of precursor in the cytoplasm. Fractionation controls showed that there was no leakage of cytoplasm into the periplasmic fraction, and the only faint mature HiPIP band in the cytoplasm indicated little proteolytic cleavage of the signal peptide or residual periplasm in the cytoplasmic fraction. The Tat deficient empty vector control strain did not show any transport and instead the bands that are indicative for cytoplasmic accumulation of HiPIP, which are a strong cytoplasmic precursor band and a band that is due to partial degradation of the signal peptide. All TatA variants with shortenings showed detectable cytoplasmic precursor bands, with only very little precursor accumulating in A-1 variants and larger quantities accumulating in A-3 variants. The A-5 variant was inactive, with no detectable mature protein in the periplasm. The TatA variants with extensions showed slightly less transport in the A+1 variant and only very little cytoplasmic precursor in the A+2, A+3, and A+4 variants, but a sudden almost complete block of transport with the A+5 variant. The A+6 variant was inactive. With the TatB variants, we observed little accumulation with the B-1 and B-3 variants, indicating only a minor defect, whereas the B+3 variant strongly accumulated HiPIP in the cytoplasm and the B+4 variant hardly transported HiPIP anymore. Interestingly, the combinations A-1/B-1, A-3/B-3, and A+3/B+3 all showed a stronger effect than the single variants, indicating that the defects were additive, or that there was indeed some partial compensation of defects of TatA by TatB or vice versa detectable in this semi-quantitative assay. Such effects were more difficult to detect by the apparently less sensitive amidase-based assays, in which few transported enzymes may often suffice for physiological complementations (compare with the SDS sensitivity assay in Fig 2A).

**Fig 3:**
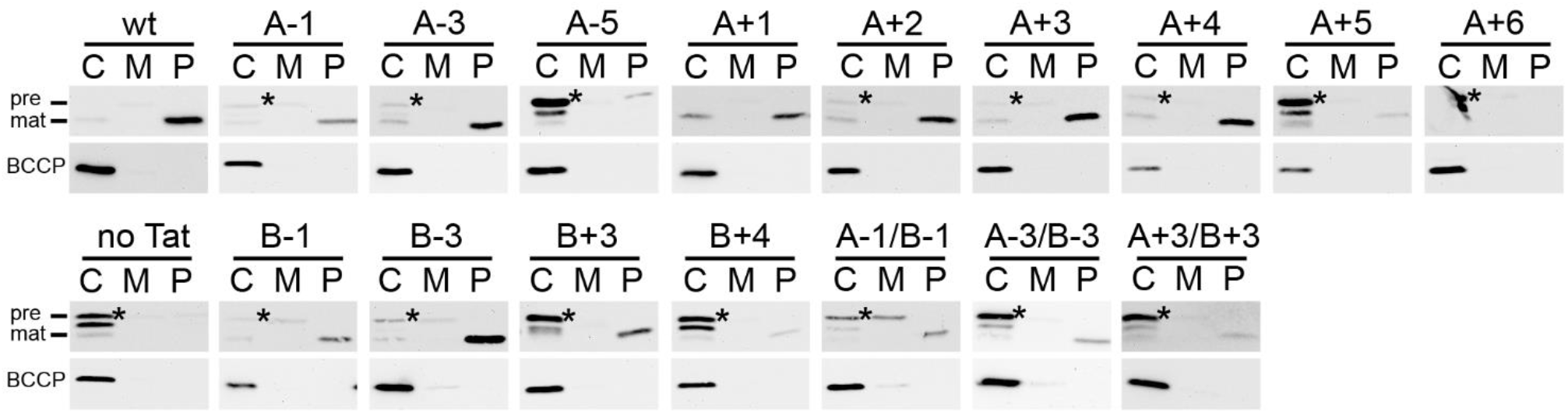
Biochemical Tat functionality assays with mutated Tat systems. Transport analysis by SDS-PAGE/Western blot detection of the precursor (pre, band at 14 kDa) and mature (mat, band at 11 kDa) forms of the Tat substrate HiPIP in subcellular fractions of cells containing the indicated single TatA or TatB variants or combinations thereof (upper blots). Biotin carboxyl carrier protein (BCCP, band at 22 kDa) was detected as fractionation control (lower blots). Cytoplasmically accumulating precursor is marked by asterisks. Note that any detection of mature protein in the periplasmic fraction indicates functional transport, whereas accumulation of precursor in the cytoplasm is only observed in cases of a partial or complete defect in transport. C, cytoplasm; M, membrane; P, periplasm.

Together, the activity measurements indicate that the 12-residues length of the hydrophobic helix in wild type TatA and TatB results in highest transport efficiency. In case of TatA, shortenings by up to three residues and extensions by up to four residues are tolerated, and some residual transport is even detectable with a five-residues extension. TatB similarly tolerates shortenings and truncations of three residues, with only little residual transport detectable in case of the four-residues extended B+4 variant.

### Variation in the TatB TMH length, not in the TatA TMH length, interferes with TatBC complex stability

It is well-described that TatB interacts with the short helix 5 of TatC, whereas TatA can interact with the similarly short helix 6 of TatC [38,28]. It was therefore possible that functional effects of length variations in the hydrophobic TMH of TatA or TatB are simply due to disturbed interactions with TatC. To examine this aspect, we analyzed the stability of the known TatC-containing complexes [27].

Membrane fractions were solubilized by digitonin and analyzed by BN-PAGE/Western blotting to detect Tat complexes TC1 and TC2 (Fig 4). While the complexes were unaffected by the length variations in TatA (Fig 4A), the B+3 and B+4 variants caused disassembly of the complexes (Fig 4B). In case of B-1, only the TC2 band was affected and changed to a very diffuse band, whereas the TC1 band remained unaffected. With B-3, the diffuse band of the size of TC2 was the only remaining complex, and TC1 had completely disappeared. Some disassembly of Tat complexes is always observed and has been described previously [15]. Together, these data clearly indicate a severe effect of the TatB variants on TatBC interaction and consequently on TatBC complex stability. As also the functional TatB variants showed this destabilizing effect, sufficient TatBC complexes must have been present in living cells to support Tat transport with this expression system.

**Fig 4:**
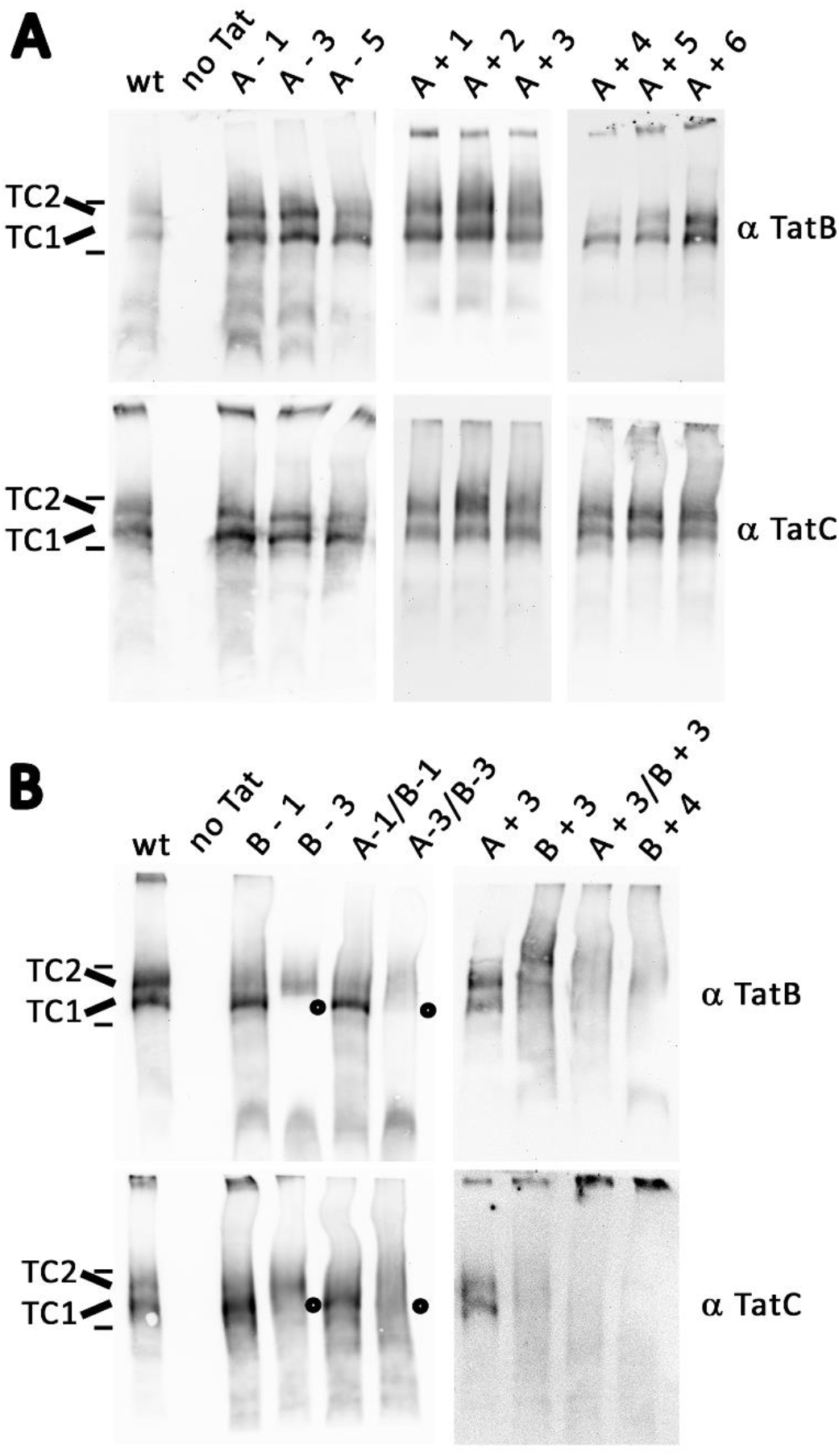
Analysis of Tat complexes in strains with mutated Tat systems. (A) BN-PAGE/Western blot detection of Tat complex 1 (TC1) and 2 (TC2) in strains containing Tat systems with indicated TatA variants (A) or with TatB variants alone or in combination with TatA variants (B). Blots were developed with antibodies against TatB (upper blots), or TatC (lower blots). The position without the TC1 band in B-3 Tat systems is highlighted by open circles. B+3 and B+4 extensions have more general destabilizing effects. For direct comparison, the A+3 system is analyzed in parallel with B+3 and the A+3/B+3 combination. Positions of TC1, TC2, and molecular weight markers (440 kDa, 669 kDa) are indicated on the left.

### METTEM-analysis of TatA clusters in the cytoplasmic membrane at wild type expression level

In the above shown BN-PAGE analyses, we could only detect effects of TatB variants on the Tat complexes, which is not surprising as most TatA dissociates from the detectable TatBC-containing complexes upon detergent solubilization, and only a regular ladder of multiple homooligomeric TatA associations is detected that does not permit analyses of the TatBC-bound TatA [33,34]. To reveal potential effects of the mutations on TatA, the lack of structural information in principle could be overcome by MD simulations. However, to create a reasonable starting point for MD simulations, we required some information about the organization of the large TatA assemblies at translocation sites. So far, it is only known from thylakoidal cross-linking analyses that at least up to 16 TatA protomers can be cross-linked at active translocation sites over a 4-minutes time span [22], which is evidence for an association of this number of protomers. Another analysis had indirectly calculated a median of 25 interacting TatA-YFP protomers in such complexes, based on mobility and fluorescence intensity measurements [23]. The pixel size of 50 nm in these analyses certainly could not resolve single protomers or even whole TatA assemblies. We thus first aimed to examine the proposed larger TatA associations in the *E. coli* system by a higher-resolution method. To our knowledge, electron microscopy is the only currently available technique that can give sufficient positional resolution within cells. We chose metal-tagging transmission electron microscopy (METTEM) for these analyses, using metallothionein-gold tags [45–50]. Metallothionein-gold tags have the advantage of directly labeling the protein of interest with rather small EM-detectable electron-dense domains. In initial experiments with abundant recombinantly produced Tat systems, we found that three metallothioneins in row (= MT3) permitted clearly visible electron-dense granules of +/- 1 nm in diameter in *E. coli* cells, and the detected particles could be identified as gold by PEELS analysis (Fig 5). Note that the recombinant protein levels that have been used for the PEELS analysis resulted in large clusters of interconnected associations that sometimes deformed the membrane, but this measurement only served to proof that the gold label of the MT3 tags had worked (Fig 5BC), and no other conclusion is drawn here.

**Fig 5:**
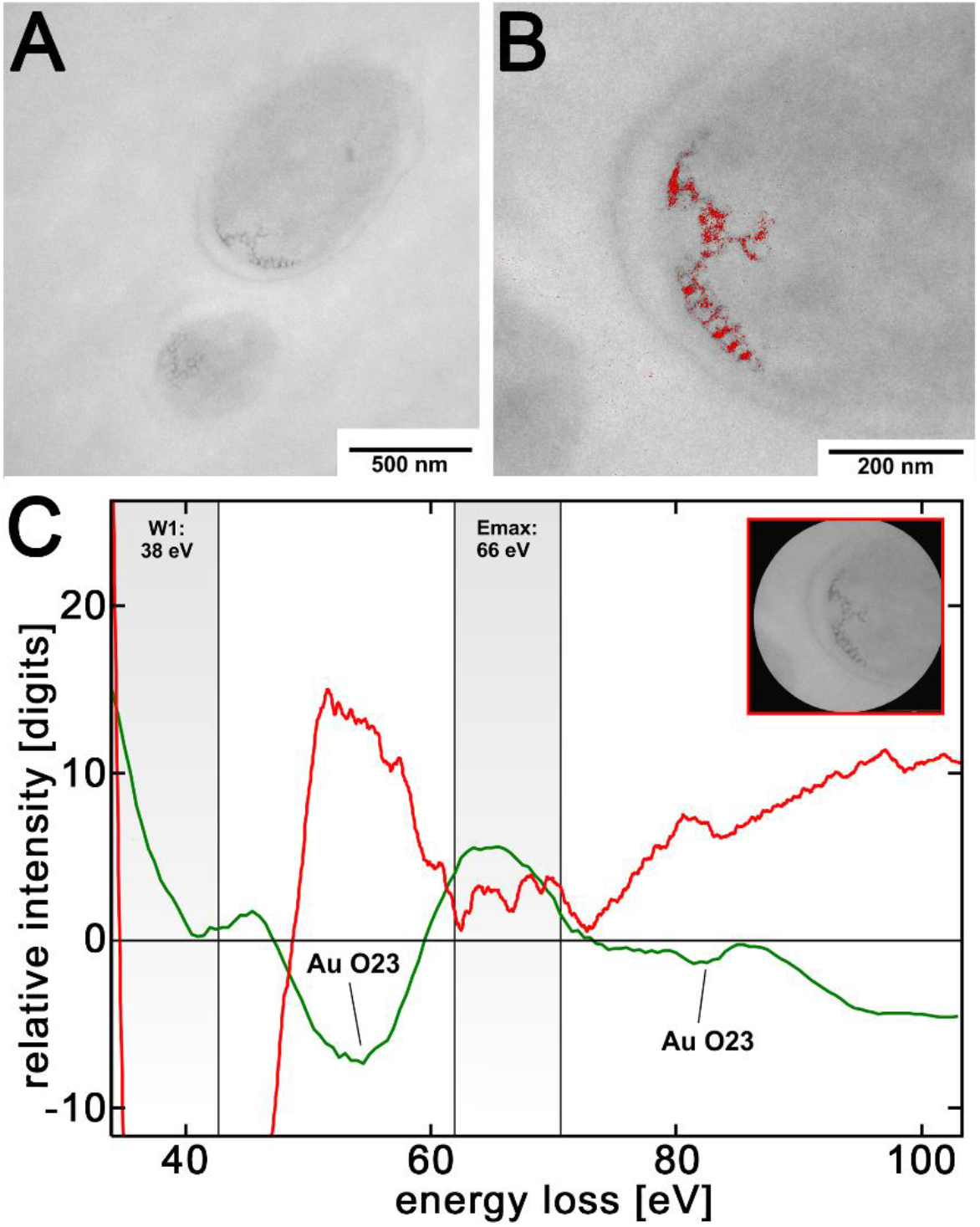
PEELS analysis of TatB-MT3-gold. (A) Survey micrograph of the analyzed cell containing the recombinant TatAB(MT3)C system as expressed by pABS-*tatAB(MT3)C*, with TatB-MT3-gold visible in clusters at the cell pole. (B) Electron spectroscopic element map at the Au-O23 edge Emax at 66 eV, energy-slit width: 8 eV, superimposed in red to the image of the cell pole shown in (A). Energy-settings and corresponding energy-slit width for the jump-ratio acquisition are indicated in C (W1; Emax). (C) PEEL-spectrum of the MT3-gold containing cell pole (inset: measuring area), detecting gold at the Au-O23 edge (red graph) and an energy maximum, which is in accordance to the Au-reference PEELS (green graph) from the EELS-atlas [51].

Having the method established, we attempted to localize the Tat components at wild type expression levels in strains that had the modified *tat* operons chromosomally integrated. With TatB-MT3 and TatC-MT3, no gold was detectable at wild type level, but it was possible to detect TatA-MT3 in these experiments, which is likely due to the higher abundance of TatA relative to TatB and TatC in *E. coli*, which are present in an estimated 50:2:1 ratio, respectively [52]. The wild type level of the TatA-MT3-abundance was confirmed by SDS-PAGE/Western blot analysis (Fig 6A). Substrates interact with the cytoplasmic face of TatA [25], and it is known that YFP tags (ca. 26 kDa) at the cytoplasmically located C-terminus of TatA inactivate TatA in the TatABC system of *E. coli* when produced at wild type level [21]. Similarly, we found that our MT3 tag (ca. 21 kDa) rendered TatA almost inactive (Fig 6B), as a slight activity was likely due to some degradation to mature size (Fig 6A). As there so far do not exist any protein tags that can be fused to TatA without compromising activity at wild type levels, this result was not surprising. Importantly, the tag did not prevent TatA-TatA interactions, and the number of TatA protomers in these associations was comparable with the number concluded from cross-linking of active non-tagged TatA (see below, 22). Therefore, the effect on activity was most likely due to steric inhibitions of substrate interactions, as substrates need to pass through the TatA assemblies. In any case, these analyses can be taken as first direct visualization of wild-type level TatA assemblies in any organism.

**Fig 6:**
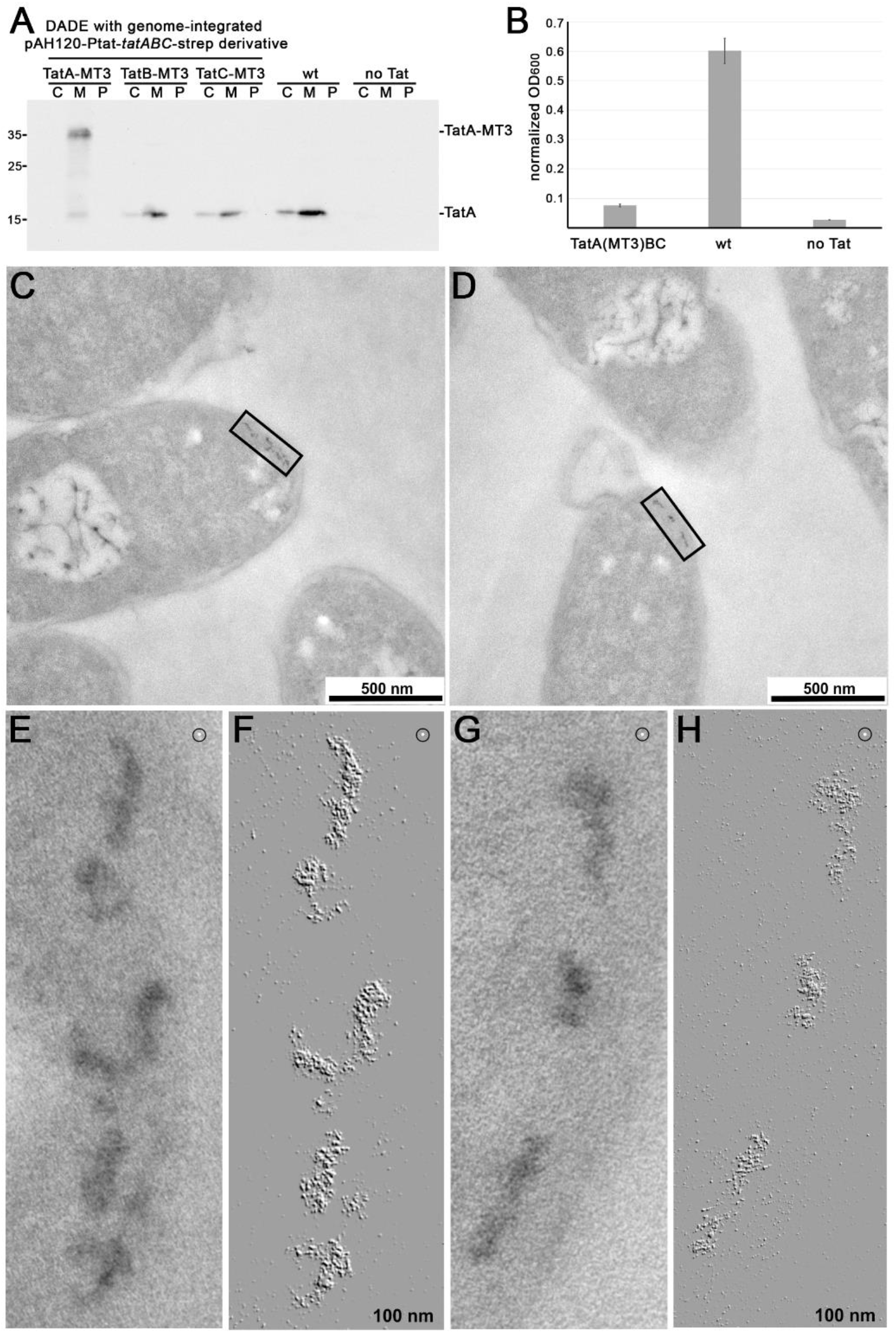
Detection of TatA-MT3-gold at wild type expression level. (A) Detection of TatA by SDS-PAGE/Western blot analysis in subcellular fractions of strains with MT3-tags at TatA, TatB, or TatC as encoded by single copy chromosomally integrated *tatABC-strep* operons. Wild type *E. coli* and the Tat-deficient strain DADE are analyzed for comparison. Positions of molecular weight markers are indicated on the left. C, cytoplasm; M, membrane; P, periplasm. (B) SDS-sensitivity measurements of strains with TatA(MT3)BC, TatABC (wt), or no Tat system. (C) and (D) show overview images of cells with gold-labeled TatA-MT3-assemblies. Areas boxed in the respective overviews are shown below at higher magnification (lower edge = 100 nm). (E) and (G) are original images, and (F) and (H) were topographically ‘relief’-filtered to discriminate MT3-gold granules in pseudo-3D within TatA-MT3 assemblies. White 1 nm-dots (encircled in E,F,G,H) have been included as a direct scale for 1nm MT3-gold-granules.

TatA-MT3 formed gold assemblies near cell poles of irregular shape when seen in quasi top-view projection and roughly normal to the electron beam (Fig 6C-H). The mean length of the gold-assemblies was 20 nm ± 6.0 nm (N = 40, min = 10 nm, max = 37 nm). The width of the assemblies was about 10 nm. To determine the number of TatA-MT3 protomers in such assemblies, we counted the number of MT3-gold particles in 35 regions containing MT3-gold, and calculated from this the average number of 17 +/- 6 MT3-gold particles per 200 nm^2^, which is the dimension of an average TatA-MT3-assembly. Interestingly, several of these assemblies visually appeared to be in contact, suggesting that they were somehow interdependent or connected (Fig 6E-H), which might be due to a clustering with TatBC complexes. Note that the MT3-gold intensity was variable, most likely due to individual kinetics of gold aggregate formation within MT3 tags, which may depend on the local environment (see supplemental Fig S1).

Sometimes, TatA-MT3-gold-assemblies appeared in longer linear arrangements in the membrane. However, tilting of the sample revealed these arrangements to result from superposition of individual assemblies when they are oriented in parallel to the electron beam within the ultrathin section (Fig 7).

**Fig 7:**
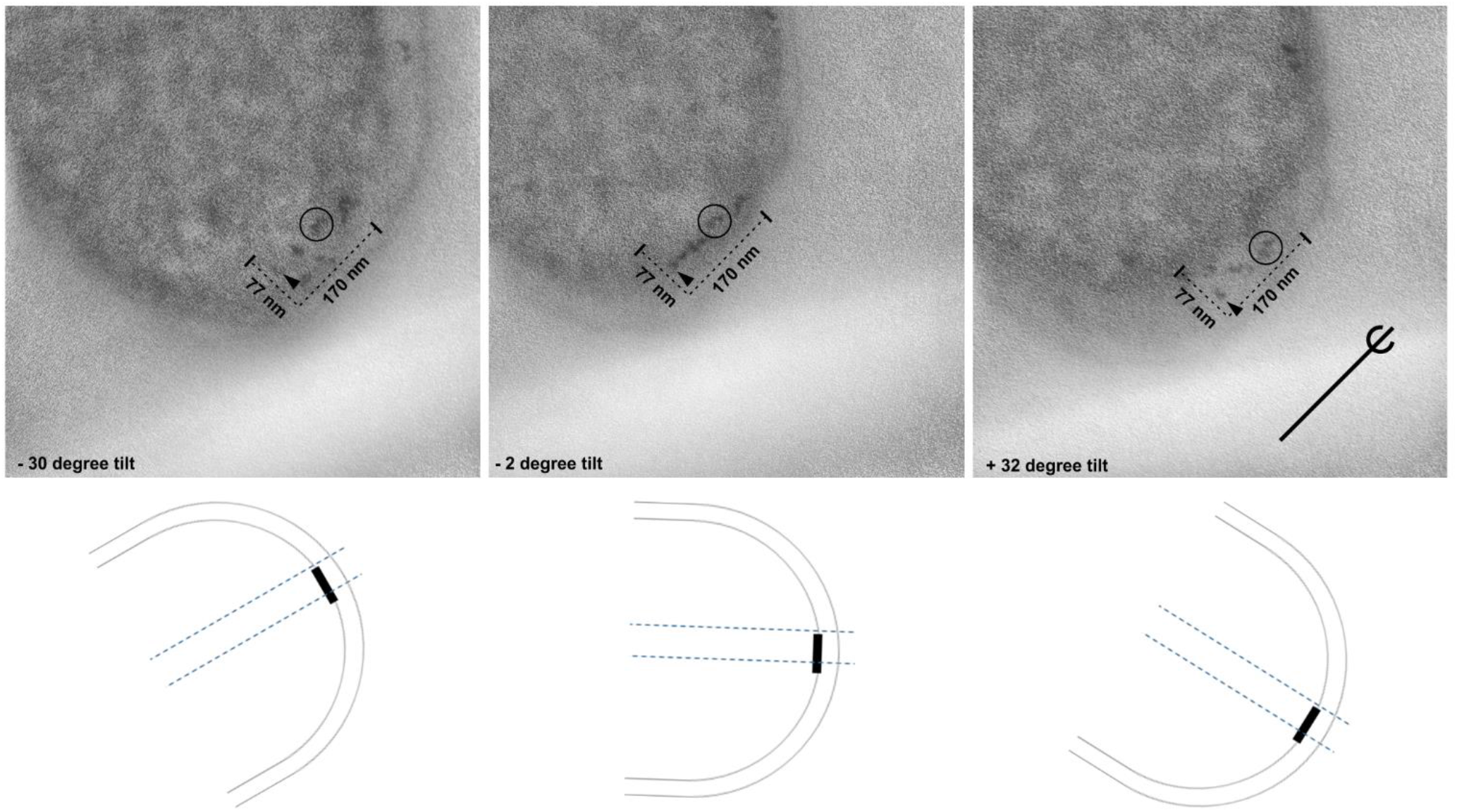
TEM-Tilting analysis of polar TatA assemblies. A seemingly very long TatA association, detectable by TEM of TatA-MT3-gold at a cell pole (middle), has been analyzed in the same cell by tilting the sample from -30° (left) to +32° (right). The tilted cell revealed that individual TatA-MT3 gold assemblies had been transformed into a single longer association due to superprojection of the assemblies when oriented parallel to the electron beam (tilt -2°).

In conclusion, TatA-MT3 formed longitudinal assemblies of irregular shape that were usually positioned near cell poles, the subcellular location of Tat translocons that has been observed in previous fluorescence-based studies [20,53]. The number of subunits in such assemblies agreed with previous calculations [23,22]. An often observed close clustering of several of these TatA assemblies suggested their indirect interaction, possibly mediated by TatBC, which explains the previous detection of a significant number of larger associations by fluorescence microscopy that was limited to 50 nm pixel resolution [23].

### MD simulation indicates membrane thinning and lipid disorder generation by the short TMHs and aligned APHs of TatA clusters

We then examined such linear TatA assemblies by 1500 ns MD simulations with 22 TatA protomers in the cytoplasmic membrane lipid bilayer, which is in the upper range of the observed TatA assemblies and therefore might reflect functional associations at Tat translocons. The MD simulations were challenged by the fact that, in living cells, larger TatA assemblies require TatBC interactions for their formation [20,21]. As no TatBC complex structure is known, and as we therefore could not include TatBC complexes in the simulations, we attempted to overcome this problem by placing linear arrangements of TatA protomers in close proximity with opposite APH-orientation. We had four reasons to expect a linear arrangement of TatA in the TatA assemblies: (1) Our METTEM analyses had detected longitudinal assemblies (see Fig 6), (2) an NMR structure of a TatA dimer showed laterally aligned APHs, which is only possible in linear arrangements [10], (3) circularly arranged TatA protomers had been reported to collapse within 40 ns in MD simulations, unless the circles were filled with a lipid bilayer that would need to be removed to permit translocation [8], and (4) a linear arrangement would permit regular interactions between TMHs of two oppositely oriented TatA protomers. This approach was successful, as the transmembrane helices readily moved into staggered positions, and the resulting complex remained stable over time in the 1500 ns simulation (Fig 8A). Most APHs laterally contacted neighboring APHs, with some interactions interrupted (Fig 8A), which agrees with the NMR evidence for lateral APH association [10], and these aligned APHs formed an angle of in average 109° with the TMH, which agrees with solid state NMR analyses on *Bacillus subtilis* TatA [54].

**Fig 8:**
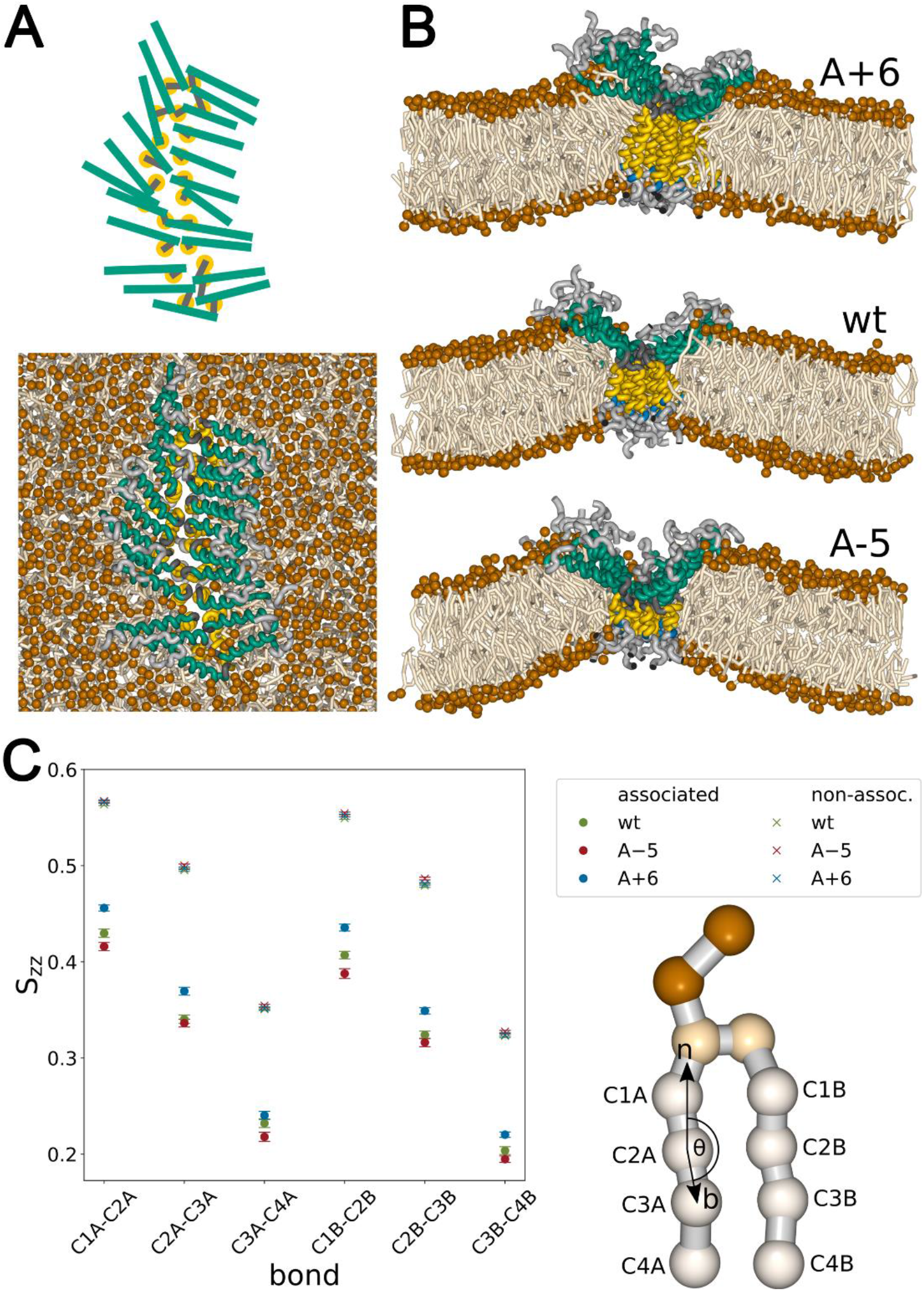
MD simulations with 22 TatA protomers of the wild type and the variants A-5 and A+6. (A) Top view from the cytoplasmic side on the 22 TatA wt simulation. (B) Cross sections through TatA complexes, with the cytoplasm above and the periplasm below the membrane cross sections. TMH, yellow; APH, cyan; Q8, turquoise; hinge region, dark grey; N- and C-termini, grey; lipids, light and dark (head groups) brown. (C) Lipid tail order parameter for lipids in the vicinity of the TMH (associated) and lipids in the bulk phase (non-associated). Bars indicate 95% confidence intervals. The shown coarse grained model of DPPE illustrates tail beads and the orientation of an example tail bond vector (b) relative to the bilayer normal (n), which is the basis for the order parameter. The other beads in the model are ethanolamine and phosphate (brown), and glycerol (light brown).

Also, the hinge was positioned deep in the membrane, which agrees with accessibility analysis data [25]. The aligned APHs formed a deep V-shaped groove at the membrane surface (Fig 8B, see S1 video). As a result of this constellation, the membrane was thinned to about half its natural thickness at the hinge region. A view on the lipids in this simulation revealed that the lipid head groups were excluded from areas of aligned APHs, and the acyl groups were therefore disordered and partly oriented in parallel to the APHs. The lipid disorder in the membranes could be quantified employing the lipid tail order parameter S_zz_ (Fig 8C). The data confirmed that the lipid order was lowered in the environment of the APHs.

We then carried out the simulations with the TatA variants A-5, A-3, A-1, A+1, A+2, A+4, and A+6. As now expected, the shortenings resulted in further thinning of the membrane, whereas the extensions of the TMH resulted in a thicker membrane, which becomes evident when comparing the simulations A-5, wt, and A+6 (Fig 8B). Accordingly, the lipid order further decreased with a shorter TMH and increased with a longer TMH (Fig 8C). The TatA simulations therefore had added a potential new aspect to the role of the short length of the TMH, as they suggested that this exact length might not only be relevant for interactions (as shown for TatB; see Fig 4) but also for membrane destabilization, which mechanistically could be important (see discussion).

As the MD simulations had indicated that large TatA assemblies can thin the membrane, we addressed whether the number of TatA protomers in the association could be relevant for membrane thinning. We repeated the simulations with 15, 10, and 5 TatA protomers, and found that the membrane-thinning effect required the lateral association of multiple APHs, which was the reason why only associations of >10 TatA protomers efficiently formed the V-shaped groove in the membrane (Fig 9AB). In case of only 5 protomers, the lateral associations of APHs were too few to impose a V-shaped groove on the membrane lipids, which is why such small associations had less effect on the membrane bilayer, as quantified by the tail order parameter (Fig 9C, see S2 video).

**Fig 9:**
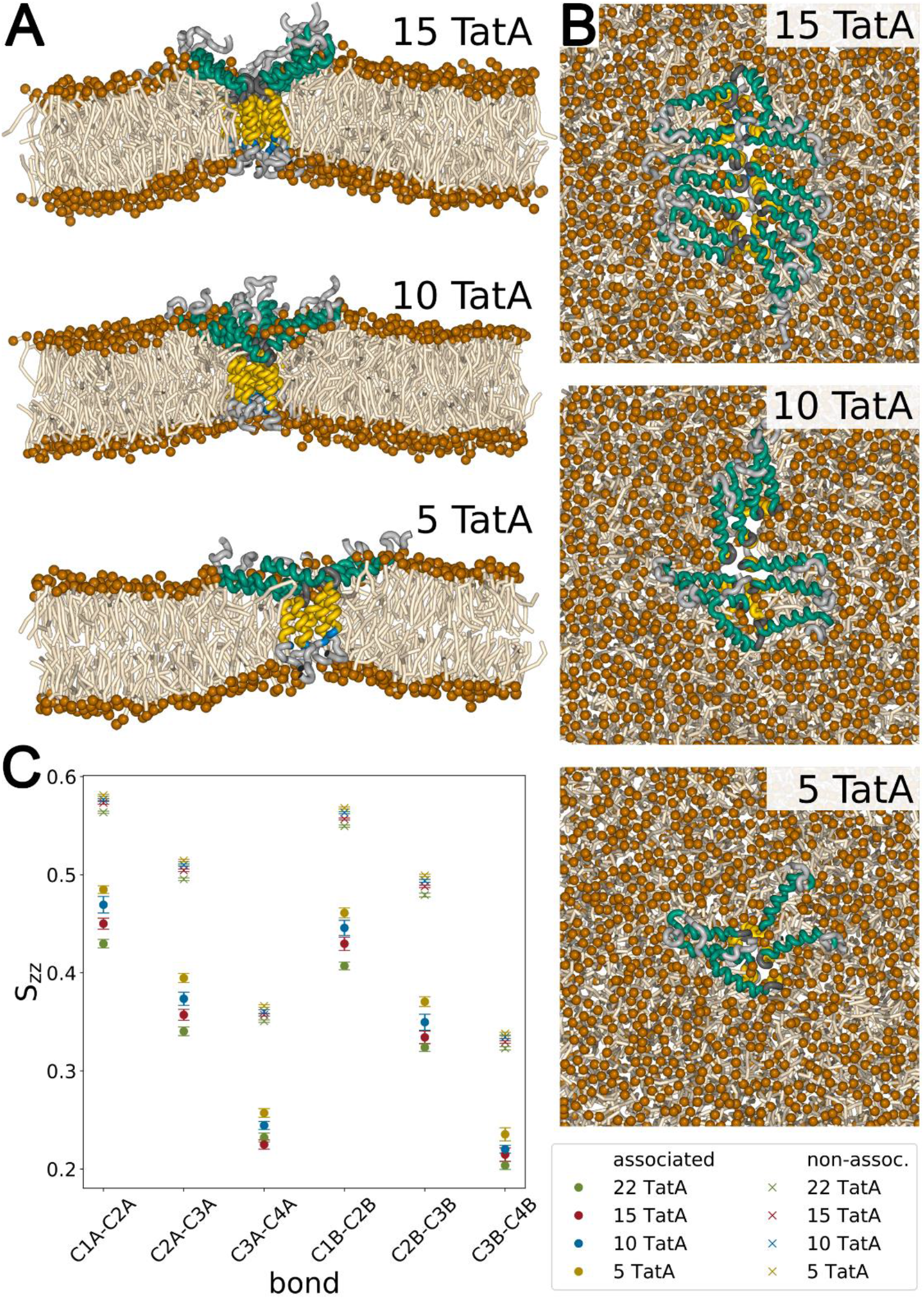
MD simulations with TatA complex of 15, 10, and 5 protomers. (A) Cross sections through TatA complexes. TMH, yellow; APH, turquoise; Q8, blue; hinge region, dark grey; N- and C-termini, grey; lipids, light and dark (head groups) brown. (B) Top views on the TatA complexes. (C) Lipid tail order parameter for lipids in the vicinity of the TMH (associated) and lipids in the bulk phase (non-associated). Bars indicate 95% confidence intervals.

### Shortening of the TatA TMH can affect cellular growth

The MD simulations suggested a second potential reason for the exact TMH length, which is the compromise between a mechanistically required membrane destabilization and an energetically required membrane stability. It has already been shown several years ago in *in vitro* experiments, that TatA can destabilize the membrane and that substrate-binding to TatA enhances this membrane destabilization, resulting in proton leakage [25]. We thus wondered whether a too short TMH could result in extreme membrane destabilization *in vivo*, which could represent a selective pressure for maintaining the minimum of 12 consecutive hydrophobic residues in TatA (and consequently also in TatB and the short helices 5 and 6 in TatC, which need to assemble with TatA at the translocation site). As proton leakage might affect growth, we analyzed growth of all strains used in our study (Fig 10).

**Fig 10:**
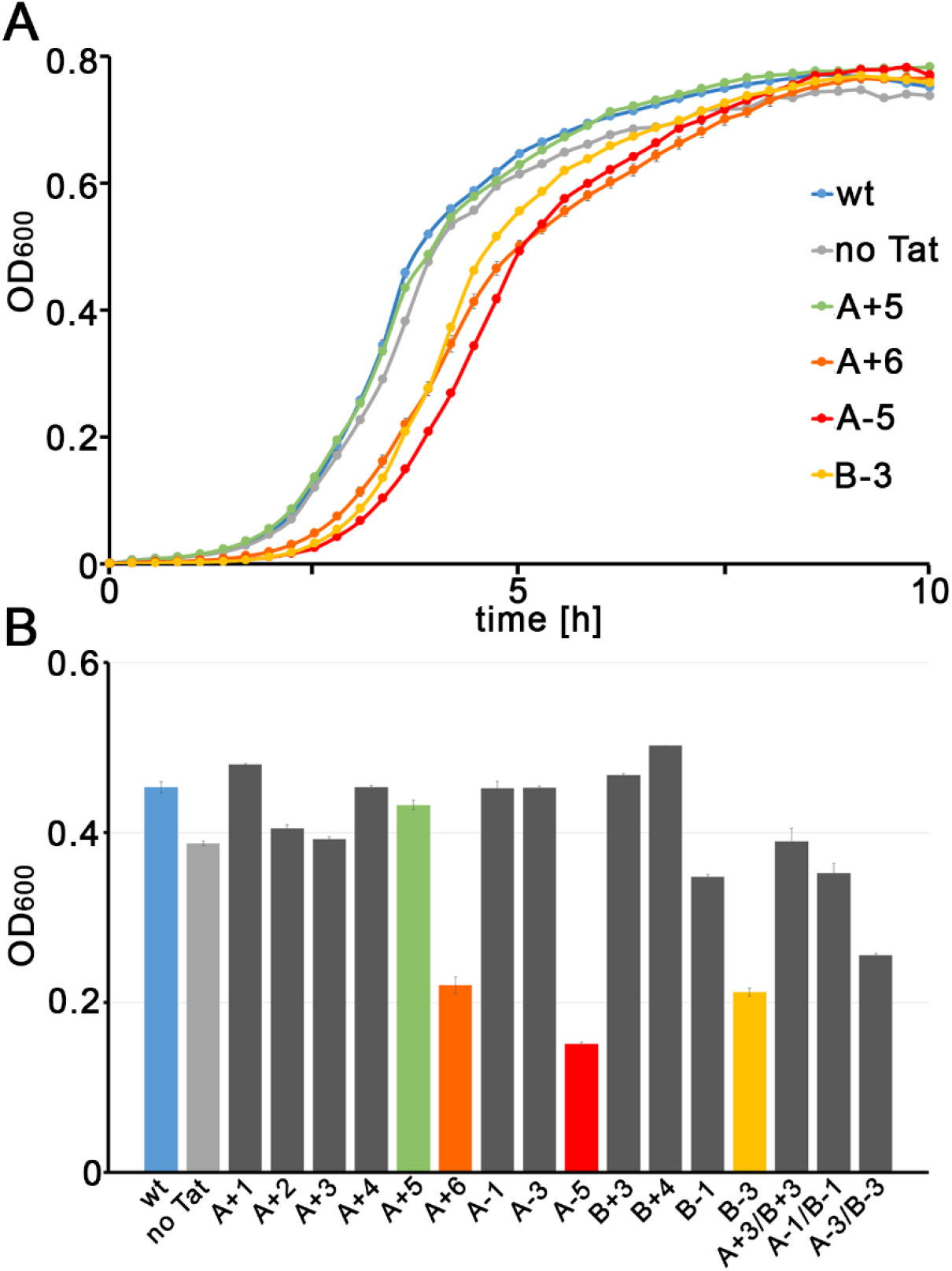
Growth analysis of strains containing the Tat systems analyzed in this study. (A) A-5, B-3, but also A+6 inhibit cellular growth. For comparison, strains with the wild type Tat system (wt) and without a Tat system (no Tat) are included, as well as the A+5 strain, which does not show a growth phenotype, although A+6 shows a clear growth phenotype. (B) Comparison of the growth phenotype of strains with all Tat systems used in this study, using the OD600 at the 3:36 h time point, as this time point is right before the end of the exponential growth of the fastest growing strains. For the strains shown in (A), the same color code is used. Error bars indicate the standard deviation of three cultures. Note that the OD600 was recorded by a 96-well multititer plate reader with 5.3 mm initial path length.

Growth curves indicated a delay of growth by A-5 and B-3, which may have been caused by cellular adaptations to stressed membranes (Fig 10). A likely cellular adaptation would be the Psp response, as Psp system components are known to stabilize membranes [55], and they have been shown to interact with Tat systems [56,57]. Interestingly, also A+6, not A+5, caused a clear growth phenotype, which may be due to proton leakage as result of an influence on structures of other membrane proteins.

## Discussion

Helices that pass the bacterial cytoplasmic membrane have to traverse a hydrophobic environment of about 3 nm [58]. However, the TMH of TatA is predicted to consist of only 15 residues [10], and that of TatB consists of only 13 residues [9], and in these helices there is only a stretch of 12 consecutive hydrophobic residues. A helix of 12 residues has a length of 1.8 nm, which does not suffice to span a lipid bilayer and therefore causes a hydrophobic mismatch. Such mismatches often have functional implications, such as the regulation of the mechanosensitive channel McsL or the aquaporin GlpF [58,59], and hydrophobic mismatches can drive membrane protein interactions [60]. This study now indicates that hydrophobic mismatches are also important for Tat functionality and Tat component interactions. The length of 12 residues is not a strict requirement for Tat transport but rather an optimal length, as little elongations and extensions of the TMH only reduce but not completely abolish function and interactions (Fig 2 and 3). In case of TatB, our data show that all length variations affect TatBC complex stability (Fig 4), and activity reductions are therefore likely due to weaker TatBC interactions. It is interesting that specific variations of the TMH in TatB can have selective effects on the different complexes, which might relate to conformational equilibria of the Tat complexes, such as previously described for mutations in TatC [27]. Observations of residual activity with extended or shortened hydrophobic stretches in TatB are suggestive for significant flexibility of interactions within TatBC complexes, which likely are not rigid but rather dynamic oligomers. Also in case of TatA, a compromised interaction with TatB and/or TatC could contribute to the observed transport defects. Interestingly, the combination of extensions or shortenings in TatA and TatB showed more severe defects than the mutations in single components, indicating that there exists some detectable functional overlap between TatA and TatB. The TatA TMH can occupy the same TatC binding site as TatB [28], which may account for this functional overlap, and as the TatC interaction is likely important also for TatA, length variation in the TMH of TatA is expected to influence activity of the system, as does the length variation in the TMH of TatB.

As BN-PAGE is not helpful to address potential effects on TatA associations, which is due to the multiple associations that TatA forms in the presence of detergent, we sought to model by MD simulations such TatA associations with wild type or mutated membrane anchors. Before doing any MD simulation, we required a more direct evidence for oligomeric TatA assemblies in *E. coli*, and the METTEM analysis served that purpose, showing that TatA-MT3 can indeed form large associations of in average 17 +/- 6 protomers, which in turn can cluster to even larger associations, suggesting bridging interactions with other components, such as TatBC. In the end, our METTEM data fully agree with the cross-linking data of the thylakoid system, in which interacting components had been trapped over minutes [22], and they are also in remarkable agreement with the mean number of 25 protomers that have been calculated from fluorescence intensity and mobility measurements of TatA-YFP foci [23]. The latter fluorescence study also reported much larger TatA-YFP associations (at least up to 100 TatA-YFP), which may correspond to the somehow interconnected TatA assemblies in our METTEM analyses. Bridged TatA assemblies likely diffuse together and therefore were not distinguishable from individual TatA assemblies within the 50 nm pixels in these fluorescence analyses [23]. In contrast, METTEM resolved the 1 nm MT3-gold tags, and therefore had a higher resolution than the current limit of ∼20 nm in super-resolution fluorescence techniques [61]. To our knowledge, this is the first application for METTEM in localization of membrane protein associations in any organism.

Having directly seen and characterized wild type level TatA associations in cells, we first simulated a larger cluster with 22 subunits, which was well in the size range of typical TatA assemblies in METTEM analysis (see supplemental Fig S1) and should be in the range of sizes of TatA associations at TatBC complexes. These simulations readily yielded stable TatA associations, with TMDs interacting with each other on a staggered position. The assemblies were rather flexible, often showing small kinks due to some irregularities in the arrangements.

Most importantly, the MD simulations revealed that a potential second function of the short TMH could be membrane destabilization. The lipid tail order parameter clearly indicates a correlation of TMH length and membrane destabilization (Fig 8C). Also important is the finding that, in these MD simulations, a larger association of TatAs is required for an efficient membrane destabilization, and that the generation of the V-shaped groove comes along with generation of membrane curvature (Fig 9). The simulations fully agree with published data: The APHs laterally interact, as previously evidenced by NMR [10], the TMHs laterally interact, as suggested by previous spin-labeling EPR studies [37], the hinge domain is positioned deep in the membrane, as supported by in fluorescence quencher accessibility studies [25], and the APHs have a tilted orientation in the membrane, as seen in solid state NMR studies with *Bacillus subtilis* TatA [54]. It is noteworthy that the average angle of the APH relative to the TMH in the simulations is near 109°, which is in good agreement with the angle determined by solid state NMR [54]. Considering the obvious mechanistic tolerance of some length variation (Figs 2 and 3), the strict conservation of the short length of 12 consecutive hydrophobic residues in TatA and TatB from bacteria, archaea, and plastids (Fig 1) is unexpected, especially if only the functionality of protein-protein interactions is assumed to generate the selective pressure for the specific length. However, if membrane destabilization is a second function of these short TMHs, which may be of fundamental importance for the transport mechanism [62], then the hydrophobic mismatch as generated by the strict 12-residue-length could be a delicate compromise between transport efficiency and proton leakage. Our MD simulations clearly show that the short TMHs and aligned tilted APHs of TatA have the potential to destabilize membranes, and the simulations of the TatA assemblies with extended or shortened TMHs show clear effects of such alterations on membrane thickness and lipid order (Fig 8). Proton-tight membranes are of key importance for energy metabolism, and bacteria have (often multiple) protective systems, which compensate for membrane destabilizations. First growth experiments already indicate that adaptation processes are required in response to A-5 and B-3 systems, and effects of minor changes in TMH length on the cellular energetization may be similar but too small to be readily detectable under the experimental conditions (Fig 9). One such adaptive system might be the Psp response system, which is known to stabilize membranes [55] and which already has been shown to interact with TatA [56]. In light of the results of this study, we would like to put forward the hypothesis that TatA, TatB and TatC together catalyze the membrane passage of folded proteins across locally destabilized membranes. The “membrane weakening and pulling” hypothesis proposed that this process could depend on membrane destabilization by multiple aligned TatA TMHs [63], which was later supported by an experimental demonstration of membrane destabilization, MD simulations and recently by electrochromic shift measurements [8,25,62]. A conformational switch of the TatA APH in response to substrate binding has been first demonstrated in thylakoids and then in the *E. coli* system, and has been suggested to contribute to this membrane destabilization [25,26].

Hydrophobic mismatches are not only generated by the TMHs of TatA and TatB, but also by the short TMHs 5 and 6 of TatC. Notably, the TMHs of TatA and TatB are both shown to interact with these two TatC helices in a resting state [24,38,28], which is likely the reason for the shown importance of the specific short length of the TatB TMH for stable TatBC associations (Fig 4). As substrate binding has been shown to trigger the association of TatA [22,64,20,21], as TatA association has been reported to precede the translocation of TatBC-bound Tat substrates [18,19], as substrate has been shown to trigger membrane permeabilization by TatA *in vitro* [25], and as TatA assemblies have the capacity to destabilize the membrane (Fig 8), we would like to suggest that larger TatA assemblies may form at TatABC complexes to enable Tat transport (see Fig 11). In principle, such an interaction would explain why most TatA is lost during purification of TatBC complexes [15], and this model also explains how a simple conformational shift of TatC helices 5 and 6 in response to substrate binding may enhance the hydrophobic mismatch at the TatBC core to induce the binding of a large number of TatA protomers to the translocation site, which induces membrane destabilization where it is needed (Fig 11).

**Fig 11:**
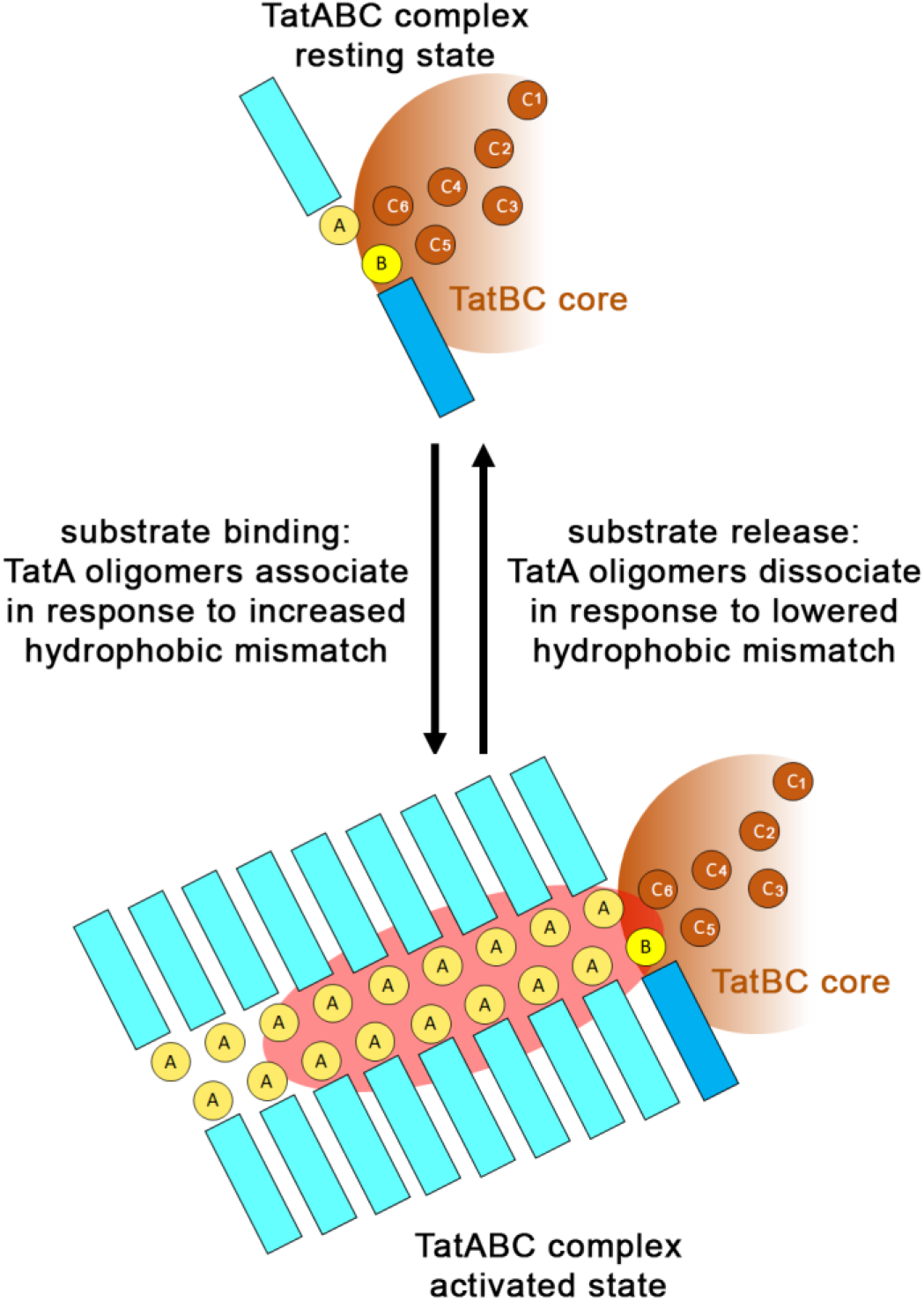
Potential substrate-induced assembly of membrane-weakening TatA-clusters at TatBC core complexes. Top view on the cytoplasmic face of the membrane, only positions of TMHs and APHs are schematically depicted. TMHs of TatA, TatB, and TatC are labeled A, B, or C, respectively. In case of TatC, which has 6 TMHs, the helix number is also indicated. The destabilized membrane region is highlighted by a reddish background. Only one docking site for TatA at the oligomeric TatBC core complex is exemplified in the scheme. The substrate that induces the conformational switch is only mentioned but not shown for clarity reasons. The color code for the TMHs and APHs is taken from Fig1. See text for details.

The model also gives an idea about the region of membrane destabilization, which is essentially the region underneath the APHs. There are two plausible pathways through the destabilized membrane: (1) The two staggered rows of TatA, which are vis-à-vis positioned in the cluster, could open and permit the passage of the Tat substrate, or (2) the TMHs remain in the center of the association and the substrate passes through the thinnest point of the destabilized membrane close to the center of the TatA cluster. As Tat substrates can be large in diameter, it is likely that both happens: TMHs may lose their contact and APHs may dissociate to permit the passage of globular proteins. In such a situation of disorder, it could well be that TatA exchanges with TatB at its TatC binding site, especially when the system is saturated with transported substrates [28]. It is also possible, that an opening of TatBC complexes can contribute to the TatA recruitment, such as proposed previously [38,24,2]. Our simulations show that TatA can form stable clusters with two interacting rows of TMHs in the center. From the energetic point of view, MD simulations indicate that there will not be any stable ring shaped TatA assembly with an aqueous hole (8). Instead, the hydrophobic mismatch can be expected to trigger associations of the TMHs, which preferentially should be linear to accommodate aligned APHs at the membrane surface that contribute to membrane destabilization (Figs 8 and 9). During the passage, transient aqueous holes would be formed that would cause uncontrolled proton flux for a very short time-span. Such a proton leakage would agree with the finding that about 80,000 protons pass through the membrane per transported protein [65], albeit proton flux does not energize the transport [66,67]. Although it is clear from these studies that the electric potential is required at certain steps of the pathway, one of which may well be the proposed conformational transition of the TatBC core complex that induces the recruitment of TatA (which is PMF-dependent; 20,21), the driving force for the transport remains to be clarified. In principle, it would be possible that the transmembrane signal peptide insertion as catalyzed by TatBC complexes [32] constitutes this driving force, as the signal peptide remains bound to the TatBC complex during transport of the mature domain [68]. Future experiments surely will shed further light on this highly fascinating transport system.

## Materials and Methods

### Bacterial strains and growth conditions

The *E. coli ΔtatABCD ΔtatE* strain DADE [42], which is a derivative of the strain MC4100 [69], was used for localization and complementation studies. *E. coli* XL1-Blue Mrf’ Kan (Stratagene, La Jolla, California, USA) or *E. coli* DH5α λ *pir*^+^ were used for cloning. Unless otherwise stated, bacteria were grown aerobically at 37°C on LB medium (1% (w/v) tryptone, 1% (w/v) NaCl, 0.5% (w/v) yeast extract) in the presence of the appropriate antibiotics (25 µg/ml chloramphenicol, 100 µg/ml ampicillin, 12.5 µg/ml tetracycline, 15 µg/ml kanamycin). For subcellular fractionations, all cultures were normalized to an OD_600_ of 1.0. For growth curves, the OD_600_ was measured in 15-min intervals with cultures grown at 37°C in shaking 96-well plates (culture volume: 200 μl), using the SpectraMax iD3 Microplate Reader (Molecular Devices, LLC., San José, California, USA).

### Genetic Methods and Plasmids

For analyses of Tat functionality and for optimization of the MT3-gold-labeling method, the TatABC components were constitutively overproduced using the natural P_*tatA*_ promoter and *tatABC* operon organization in plasmid pABS-*tatABC* and derivatives thereof [41]. The shortenings and extensions of the hydrophobic helices in TatA and TatB were carried out by QuikChange mutagenesis. Primers are listed in Table 1.

**Table 1:**
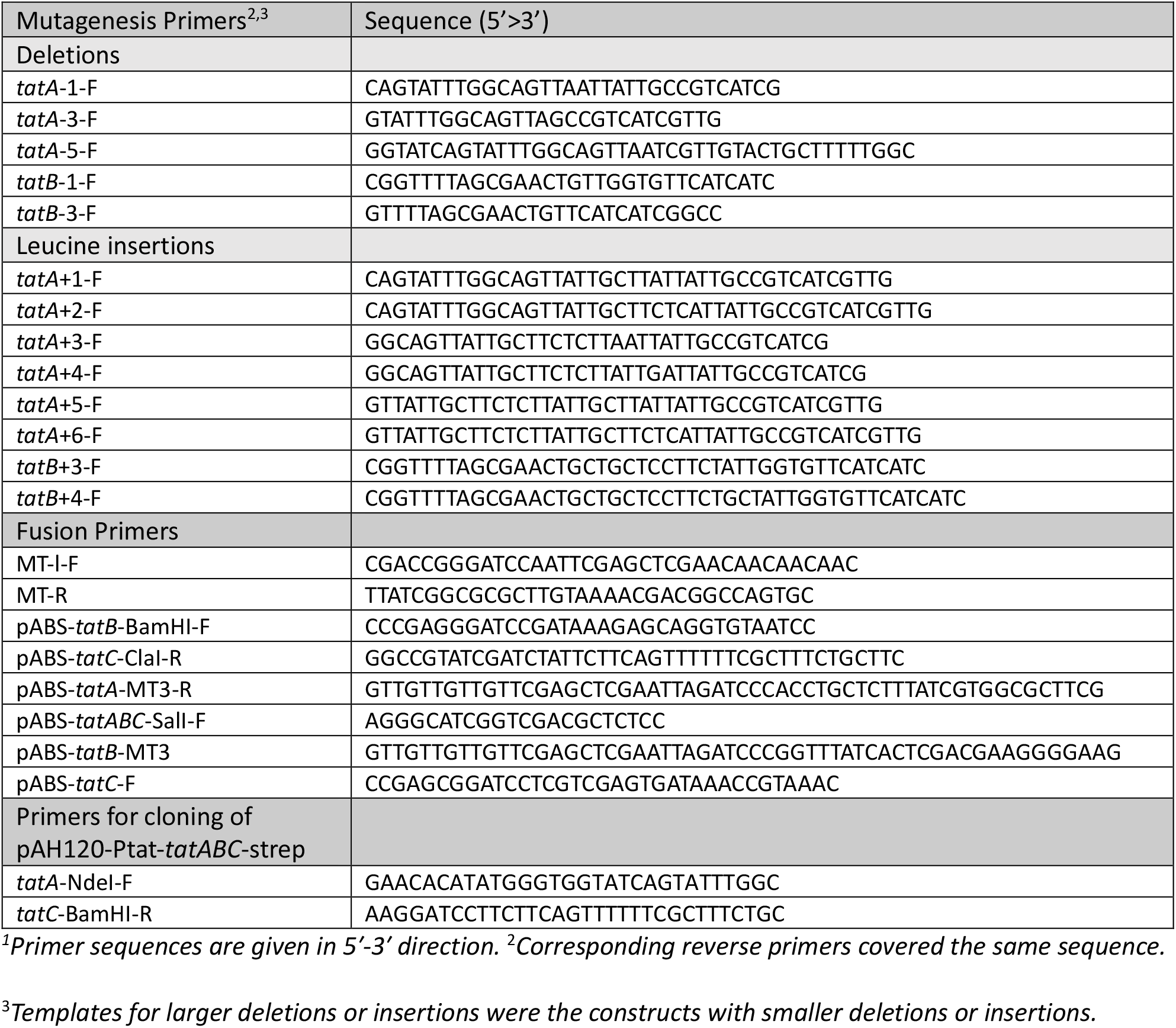
Primers used in this study^1^.

To analyze MT3-gold labeled Tat components at wild type level, the *tatABC* operon and its derivatives were cloned into pAH120-Ptat-*tatABC*-strep (see primer table), which is a *tatABC*-containing derivative of pAH120-Ptat*-tatA*-strep-ΔBamHI [70]. Briefly, the MT3-tag (including the linker) was amplified from the pMAL-c2x-MT3 [50] using primers MT-l-F and MT-R, cut with BglII/BssHII, and ligated into the backbone of the plasmid pAH120-Ptat-*tatABC*-strep cut with BamHI/BssHII. This resulted in pAH120-Ptat*-tatABC*-MT3, which was then cut with NdeI/BamHI and ligated into the corresponding sites of pABS-*tatABC*-H6 [27], resulting in pABS-*tatABC*-MT3. To construct pABS-*tatA*-MT3, *tatA* was amplified from pABS-*tatABC*-MT3 using primers pABS-*tatA*-MT3-R and pABS-*tatABC*-SalI-F, the fragment was cut with SalI/SacI and ligated into the plasmid backbone obtained by restriction of pABS-*tatABC*-H6 with the same enzymes. The same procedure was followed to construct pABS-*tatAB*-MT3, except that the primers utilized were pABS-*tatB*-MT3-R and pABS-*tatABC*-SalI-F. To append the *tatBC*-genes to the *tatA*-MT3-construct, these were amplified from the pABS-Ptat-*tatABC*-MT3 plasmid using primers pABS-*tatB*-BamHI-F and pABS-*tatC*-ClaI-R, the fragment was cut with BamHI/ClaI and ligated into the respective sites of the pABS-Ptat-*tatA*-MT3 backbone. The same steps were taken to construct the pABS-*tatAB*-MT3-*tatC* plasmid, only that primers pABS-*tatC*-F and pABS-*tatC*-ClaI-R were used and the fragment was ligated into the backbone of pABS-Ptat-*tatAB*-MT3. Finally, both the *tatA*-MT3-BC and *tatAB*-MT3-*tatC* constructs were transferred back into pAH120 by restriction of pAH120-Ptat-*tatABC*-MT3, pABS-*tatA*-MT3-*tatBC* and pABS-*tatAB*-MT3-*tatC* with NdeI/XbaI, and ligation of the *tat*-fragments into the pAH120-backbone. Vectors were integrated into the chromosomal lambda attachment site according to the protocol of Haldiman and Wanner [71]. pRK-*hip* was used for constitutive low-level expression of the *hip* gene from its own promotor [44].

### Biochemical methods

Subcellular fractionations into periplasm, membranes, and cytoplasm were performed with 50 mL cultures as described previously [70]. SDS-PAGE analysis was carried out by standard methods [72]. BN-PAGE was performed as described previously [34], but without *β*-mercaptoethanol in the buffers. TatABC complexes were solubilized with 1% digitonin. For immunoblots, proteins were semi-dry blotted on nitrocellulose membranes and blots were developed using antibodies directed against synthetic C-terminal peptides of TatA, TatB, TatC or polyclonal rabbit serum against purified HiPIP, using the ECL system (GE Healthcare, Solingen, Germany) for signal detection. Horseradish peroxidase (HRP)-conjugated goat anti rabbit antibody (Roth, Karlsruhe, Germany) served as secondary antibody. Biotin carboxyl carrier protein (BCCP) was detected by horseradish peroxidase-coupled Strep-Tactin (IBA, Göttingen, Germany). The chain formation phenotype was assessed by phase contrast microscopy. SDS sensitivity was determined by aerobic growth in LB medium containing 4% (w/v) sodium dodecyl sulfate (SDS), using the quotient of the OD_600_ with/without SDS after 3 h of growth [43]. For carbonate washes of membranes, samples were treated as described before [25].

### Electron microscopy – Sample preparation

Bacterial cells (20 ml culture, 5 % inoculum) were grown at 37°C at 140 rpm to OD_600_ ∼0.5, then 0.4 ml of an aqueous 25 mM tetrachloroauric acid solution were added, and growth was continued for 90 min at 37°C and 120 rpm. Cells were sedimented (10 min, 3,820 x g, 4°C), resuspended in 1.5 ml 100 mM HEPES pH 7.0, 90 mM sucrose (buffer A) and incubated for 5 min at room temperature (RT). Cells were sedimented again (2 min, RT, 6,800 x g) and the incubation/resuspension was repeated. Cells were then resuspended in 1 ml buffer A containing 2.5 % glutaraldehyde, incubated for > 20 min at RT, washed with 1 ml buffer A, sedimented (2 min, 6,177 x g, RT), and washed twice in 100 mM HEPES pH 6.9, 90 mM sucrose, 10 mM MgCl_2_, 10 mM CaCl_2_, sedimented again and embedded either in SPURR epoxy resin [73] or in LR-White resin [74]. 70 nm ultrathin sections were cut using an ultramicrotome with a diamond knife (Ultracut; Leica Biosystems, Nussloch, Germany) and picked up with neoprene-coated Cu-grids (300 mesh, hexagonal). *TEM image registration and processing -* Unstained samples were analyzed in the elastic bright-field mode with an in-column energy-filter transmission electron microscope (Libra 120 plus, Zeiss, Oberkochen, Germany). The energy-selecting slit aperture was set to 15 eV energy-width, and images were recorded with a cooled bottom-mount 2 × 2 k CCD (SharpEye, Tröndle, Wiesenmoor, Germany) at nominal magnifications from × 12,500 to × 50,000 near Gaussian or at 500 to 1,500 nm underfocus. Data acquisition and spectrum/image-analysis were done using the iTEM software package (Olympus Soft Imaging Solutions Ltd.). Images were further processed with the Corel Draw Graphics Suite X7 (Corel Ltd.) and ImageJ Fiji software [75]. *Parallel Electron Energy-Loss Spectroscopy (PEELS) -* PEELS was done with unstained 35 nm ultrathin sections of epoxy resin-embedded *E. coli* strain DADE/pABS-*tatAB(MT3)-*C with MT3-gold as the only electron dense intracellular material. PEELS registration was performed in the energy range from 30 to 139 eV, specific for the Au-O_23_ edge, with a 100 µm spectrum entrance aperture and 5 acquisition cycles pro record. Spectrum magnification was set to 200-fold, and the illumination aperture to 0.16 mrad with 4 µA emission current; exposure time: 0.2 s, nominal magnification: x 31,500. *Electron spectroscopic element mapping* Data acquisition was performed on well-defined Au-MT3-clusters and the energy-selective slit was set to 8 eV width. Jump-ratio images were registered at 66 eV (= Emax) and 38 eV (= 1^st^ window). These energy-settings were taken from PEEL spectrum data of the MT3-gold assemblies of interest. Exposure time: 2 s, illumination aperture: 0.63 mrad, nominal magnification: x 31,500.

### MD simulation

Molecular dynamics simulations were performed using the GROMACS simulation package, version 2019.5 [76]. The MARTINI v2.1 coarse-grained force field [77–79] was applied to simulate lipids, peptides, and solvent. In all simulations, the system was coupled to a constant temperature bath by v-rescale algorithm [80] with a relaxation time of 1.0 ps. Simulations were performed at 320 K. Periodic boundary conditions were applied to simulate bulk behavior. The time step used in the simulation was 20 fs. The dielectric constant in the simulations was ε_r_ = 15. The pressure was coupled to 1 bar employing the Parrinello-Rahman barostat [81] with a relaxation time of 12.0 ps. Unless stated otherwise, the TatA membrane system consisted of 22 TatA proteins in a lipid bilayer with 75 % phosphatidylethanolamine, 19 % phosphatidylglycerol and 6 % cardiolipin, which is a usual ratio of these lipids in the cytoplasmic membrane of *E. coli* [82]. For the MD simulations of TatA associations, a coarse-grained TatA model up to residue Glu47 was constructed based on the known NMR structure (PDB 2MN7, 10). To model the linear assembly of TatA proteins observed in the TEM experiments, 22 copies of the equilibrated TatA structure were placed vis-à-vis in two rows of 11 subunits, and inserted in the membrane via deletion of all overlapping membrane lipids. The overall dimension of these 22-mers, about 20 nm in length, closely match to the ones observed in METTEM. This system was then equilibrated for 500 ns to enable adaption of the membrane. The TMHs readily associated in a staggered way with all TatA proteins included in the linear complex. For all simulations, the TatA sequences up to position Lys49 were used, as C-terminal truncation analyses have shown that a TatA of exactly that length is still functional [12]. MODELLER 9.24 [83] was used to create the TatA variants. The resulting data were evaluated by MDAnalysis 0.20.1 [84]. The lipid tail order parameter S_zz_= ½ (3(**b**\***n**)^2^-1) was determined to quantify the average orientation of tail bond vectors **b** with respect to the bilayer normal **n** [85]. Order parameter time series were checked for equilibration and decorrelated using the pymbar package [86–88]. Angles between the TMH and the APH were calculated between two vectors connecting the center of masses (COM) of residues Gly21, Thr22, and Lys23 for the hinge either with the COM of residues Ala13, Val14, and Ile15 for the TMH, or with the COM of residues Ile28, Gly29, and Ser30 for the APH (wild type numbering). Images of the simulations were generated using NGLview v2.7.7 [89].

## Acknowledgements

We thank Sybille Traupe and Katrin Gunka (LUH, Hannover) for excellent technical assistance. We gratefully acknowledge the skillful work and experience in electron microscopic sample preparation by Inge Kristen, Sabine Schmidt and Ina Schleicher (HZI, Braunschweig).

## Supporting information

**S1 Fig.**
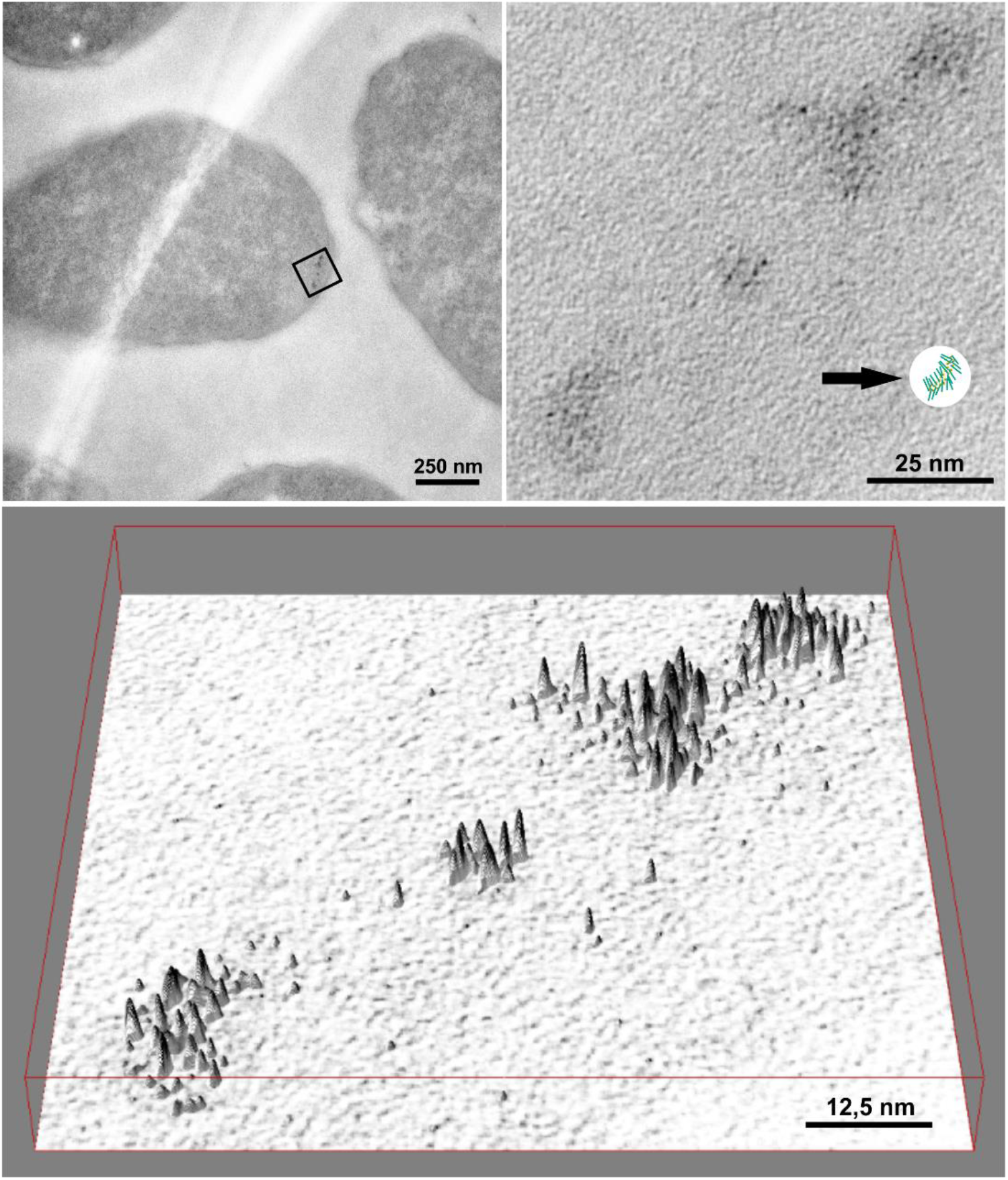
Detailed qualitative aspects of TatA-MT3-gold labeled assemblies in *E. coli* at wildtype level. (A) Survey view of an individual cell with a TatA-MT3 cluster, labeled with electron-dense gold (squared frame) in the subpolar region. (B) Detailed view of the area framed in (A), showing the TatA-MT3 assemblies of the cluster at higher magnification. Note the electron-dense granules of individual TatA-MT3-gold tags, which are about 1 – 1.5 nm in size. An isoscale scheme of the MD-simulated TatA assembly of 22 protomers (arrow, see Fig 8A) is included for comparison. Note that the MT3 domains are 69 residues apart from the modeled APHs (green). (C) A tilted, 3D-surface plot shows the range of gray level variance above the cytoplasmic background within the assemblies, which is indicative for MT3-gold. High peaks with black caps represent gold granules, as they can be readily recognized in (B). The range of different peak heights from high to low as seen in (C) indicates a variability of gold-loading of MT3 tags during the incubation of the cells with tetrachloroauric acid, possibly caused by differences in the local environment and unequal accessibility of gold.

**S1 Video. Cross-section video of the 1.500 ns simulation of a TatA-assembly consisting of 25 protomers**. Note the V-shaped groove formed by the association of the short TMHs in conjunction with the aligned APHs.

**S2 Video. Cross-section video of the 1.500 ns simulation of a TatA-assembly consisting of 5 protomers**. Note that 5 TatA protomers do not suffice to significantly thin the membrane.

